# A Geometrical Class of Population Growth Models, across Scales and Population Structures

**DOI:** 10.64898/2026.10.09.757927

**Authors:** Thomas Tunstall, Alessio Zaccone

## Abstract

We develop a general geometrical framework for growth processes in which the growth rate depends exponentially on the number of active growth-supporting components. The framework separates the fraction of active components from the size of the accessible component space, allowing qualitatively different growth laws to arise from the same underlying construction. Bacterial population growth provides an example of a fixed-size component space controlled primarily by activation and deactivation, whereas examples such as human population growth or speciation provide an expanding component space in which interactions between individuals promote further growth. This motivates a Replicative geometrical law, *dX/dt* = *αX* exp(*βX*^*γ*^), whose exponent characterizes the scaling of the effective growth-supporting component space and is tested against historical human-population regimes and eukaryotic lineage-through-time data. For innovation indicators such as patents and arXiv manuscript submissions, where an existing item need not itself reproduce, the same construction motivates a Direct Geometrical production law without the prefactor *X*. The framework therefore provides a common description of fixed and expanding component spaces while distinguishing Replicative from Direct Geometrical growth.

## I. INTRODUCTION

Models of population growth range from phenomenological fitting laws to mechanistically derived descriptions that attempt to connect macroscopic growth to microscopic processes. This distinction is particularly evident in bacterial growth, where logistic and related functions are frequently used for empirical fitting [1–4], while more detailed approaches seek to connect growth to the physiological state of the cell. In particular, bacterial growth rate is closely connected to nutrient availability, macromolecular synthesis, and cellular resource allocation, with the nucleotide alarmone (p)ppGpp providing one of the best-characterized mechanisms for coordinating environmental conditions with growth [5].

A recently proposed geometrical model [6] provides an intermediate description. Rather than attempting to model every biochemical process explicitly, it uses statistical and geometrical arguments to describe how the inactivity of growth-supporting “components” of a cell - which may represent organelles, biochemical processes, efflux systems, nutrient availability, or other effective degrees of freedom - reduces the growth rate from its maximal value. The essential idea is that growth depends not only on the existence of these components, but also on the fraction of them that are active at a given time.

Here we explore the extent to which this geometrical construction can be generalized to growth processes occurring at very different scales and population structures. The central distinction that emerges is between systems in which the accessible component space is essentially fixed, systems in which it increases but its effect on growth is biologically bounded, and systems in which newly generated components themselves enlarge the space of possibilities available for subsequent growth without per-capita growth.

This distinction provides a natural connection with the nonlinear rate-feedback equation introduced by Trachenko and Zaccone [7], in which the per-capita growth rate depends exponentially on the population variable. Rather than interpreting such exponential feedback as an indefinitely increasing microscopic reproductive rate, we show that it arises naturally in systems with an expanding configuration space, such as technological innovation and, potentially, biological diversification.

We first formulate the general geometrical framework and recover bacterial population growth as a fixed-component-space realization. We then consider expanding component spaces through human population growth, for which existing entities reproduce and the geometrical factor modulates a per-capita rate. We examine historical human-population regimes within the generalized Replicative framework and discuss how the fitted exponent and the sign of the effective feedback vary between regimes. We also discuss the applicability of this model to speciation, as speciation is also modulated by the presence of other species in a similar way. We next distinguish this Replicative Geometric model from a Direct Geometrical production law appropriate to innovation indicators that need not reproduce on a per-capita basis.

## II. GENERAL GEOMETRICAL FRAMEWORK

Consider a growing system containing an effective set of *M* (*t*) growth-supporting components. Let *A*(*t*) denote the fraction of these components that are active, with

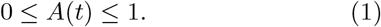

The number of active components is therefore *M* (*t*)*A*(*t*). Components need not contribute directly to growth; instead, they may support downstream processes which themselves facilitate growth. If a component is rendered inactive, all processes that depend upon it may be affected. Consequently, the influence of inactive components need not be additive.

The geometrical construction of Ref. [6] maps this problem onto overlapping excluded regions in an abstract configuration space [8, 9]. The remaining accessible volume corresponds to the effective productivity of the system. The resulting per-capita growth rate,

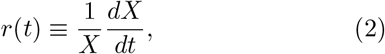

is taken to depend exponentially on the number of active components,

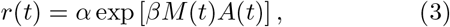

where *α* sets the baseline growth-rate scale and *β* quantifies the contribution of each effective active component. Equation 3 can also be motivated independently of the geometrical construction. Suppose that each active component multiplies the per-capita growth rate by a factor 1 + *ϵ*. Then

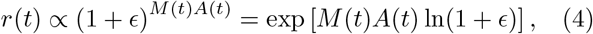

which is equivalent to Eq. 3 after redefinition of *β*.

An immediate experimental diagnostic follows from Eq. 3 when an independent physiological proxy for *A*(*t*) is available. For constant *M* = *M*_0_,

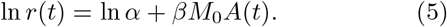

Thus, if *A*(*t*) can be inferred independently from the activation-deactivation dynamics, the geometrical model predicts a linear relationship between ln[(1*/X*)*dX/dt*] and *A*(*t*).

The physical content of the model is therefore determined by the dynamics of two distinct quantities: the fraction *A*(*t*) of active components and the size *M* (*t*) of the accessible component space.

## III. FIXED-SIZE COMPONENT SPACE: BACTERIAL POPULATIONS

In recent work [6, 10], populations of *P. aeruginosa* and *S. aureus* were studied in monoculture and co-culture, in both open and closed environments. Here we focus on the closed-system case for simplicity.

At the level of an individual bacterial cell, the effective components may correspond to metabolic processes, transport mechanisms, nutrient availability, macromolecular synthesis, or other physiological degrees of freedom. These degrees of freedom should not be interpreted as a set of literally independent molecular switches. Bacterial growth emerges from coupled physiological and regulatory processes linking nutrient availability, macromolecular synthesis, resource allocation, cell size, and cell-cycle progression. In particular, (p)ppGpp provides a well-characterized regulatory mechanism through which cellular supply and demand are coordinated with growth [5]. Homeostatic growth control is nevertheless cell autonomous, and substantial growth-rate heterogeneity can therefore exist among individual cells within the same population [5].

Within the present coarse-grained framework, *A*(*t*) should consequently be interpreted as an effective fraction of growth-supporting physiological degrees of freedom that are available or active at a given time. The assumption of independent activation or deactivation introduced below is a statistical assumption at the level of these effective components, rather than a statement that the underlying molecular and physiological processes are microscopically independent.

The total accessible component space is taken to be approximately fixed on the timescale of population growth, so that

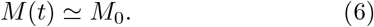

The nontrivial dynamics therefore arises primarily from changes in the effective active fraction *A*(*t*).

A component can be rendered inactive by nutrient limitation, toxins, environmental stress, or other perturbations. Conversely, components can become reactivated when favourable conditions are restored. If activation events are statistically independent at the level of these effective components, then following the transfer of a starved cell to a nutrient-rich environment one obtains

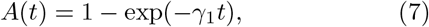

which produces the initial sub-exponential growth associated with the lag phase.

When nutrients later become limiting, components deactivate independently. Introducing the inactive fraction *I*(*t*) = 1 ™ *A*(*t*) and applying the affine transformation required for the growth rate to vanish when all relevant components are inactive, the model of Ref. [6] becomes

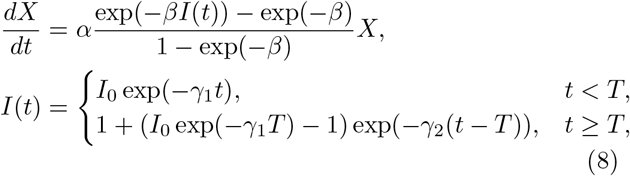

where *T* is the time at which nutrient limitation becomes important, while *γ*_1_ and *γ*_2_ are the activation and deactivation rates, respectively. In [6] an affine transformation is applied to the growth rate to ensure that cell growth tends to 0 as *I*(*t*)→1.

The bacterial case therefore represents the simplest realization of Eq. 3: the size of the accessible component space is approximately constant, while growth dynamics is generated by temporal changes in the active fraction.

### IV. POPULATION-DEPENDENT COMPONENT SPACES: HUMAN POPULATIONS

A qualitatively different regime arises when newly generated components themselves become building blocks for subsequent growth. In this case, not only the active fraction but also the space of possible future components can expand dynamically.

Human populations form highly connected social and technological networks. The survival and reproductive success of an individual depend on goods, services, infrastructure, knowledge, medicine, energy, and other resources produced by many other individuals. We can imagine this graphically, where each human is a node and directed edges between nodes represent the output of their labour which facilitates growth and reproductive fitness. These edges take the place of growth-facilitating components: these interactions define an effective space of growth-supporting components whose size we allow to scale as

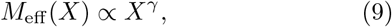

*M*_eff_ (*X*) *∝ X*^*γ*^, (9) with *γ >* 0. For a simple network in which components correspond strictly to distinct links per individual, one expects *γ ≤* 1 [11]. More generally, however, the effective components may include the outputs of interactions, higher-order collaborative structures, infrastructure, and other collectively generated resources. In this broader interpretation, superlinear values *γ >* 1 are not excluded a priori and can be determined empirically. Absorbing the proportionality constant in *M*_eff_ (*X*), together with any approximately constant active fraction, into the effective parameter *β*, the ordinary replicative factor *dX/dt ~ X* is therefore modulated by the geometrical rate factor exp(*βX*^*γ*^). This gives the human population growth law:

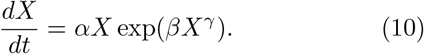

Note that *γ* = 0 corresponds to a traditional exponential growth law, whereas *γ* = 1 corresponds to the Trachenko-Zaccone growth law [7]. In this way, Eq. 10 can be interpreted as a component-space extension of the Trachenko-Zaccone model, or a ‘generalised T-Z’ model. In the narrower context of this describing a system where replicating individuals drive the growth of other individuals, we shall term this a Replicative Geometric growth model.

To test this interpretation without imposing the original *γ* = 1 limit, we reanalyse the empirical global-population data tabulated by Sojecka and Drozd-Rzoska [12], which also underlie the population analysis of Ref. [7]. Within each historical window we define a regime-wise normalized population

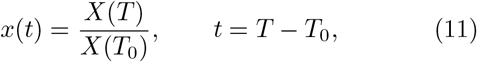

so that *x*(0) = 1, and fit

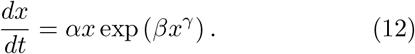

For comparison, the same data are fitted to ordinary exponential growth, *dx/dt* = *αx*, and to a power law, *dx/dt* = *αx*^*p*^: see Fig. 1 and Table I. Residuals are evaluated in ln *x*, consistently across the competing models. The three historical intervals display distinct behaviours of the same generalized construction, as summarized in Fig. 1. For 1790–1948, the unconstrained optimization yields a best-fit exponent *γ ≃*2.2, with *α ≃* 3.8 *×*10^ℒ3^ yr^ℒ1^ and *β≃* 0.16. Within the effective-component interpretation introduced above, this corresponds to a superlinear scaling of the fitted growth-supporting component space.

**TABLE 1.** The values of the parameters of the Replicative Geometric, power-law, and exponential models as fitted to human population data [12], to 2 significant figures, across three time windows: 1790 ℒ 1948, 1930ℒ 1978 (centre), and 1970 ℒ 2023 (right). For each model the AIC and BIC values are reported, to 2 significant figures.

| Model | CEF 1790–1948 |  |  |  |  | CEF 1930–1978 |  |  |  |  | SEF 1970–2023 |  |  |  |  |
| --- | --- | --- | --- | --- | --- | --- | --- | --- | --- | --- | --- | --- | --- | --- | --- |
| | $\alpha$ | $\beta$ | $\gamma$ | AIC | BIC | $\alpha$ | $\beta$ | $\gamma$ | AIC | BIC | $\alpha$ | $\beta$ | $\gamma$ | AIC | BIC |
| Exponential | $5.5 \times 10^{-3}$ | - | - | -350 | -350 | 0.013 | - | - | -220 | -220 | 0.016 | - | - | -360 | -360 |
| Replicative Geometric | $3.8 \times 10^{-3}$ | 0.16 | 2.2 | -430 | -430 | $1.3 \times 10^{-89}$ | 200 | $7.7 \times 10^{-3}$ | -300 | -290 | 0.037 | -0.57 | 1.0 | -620 | -620 |
| Power-Law | $4.0 \times 10^{-3}$ | - | 2.0 | -420 | -420 | $9.3 \times 10^{-3}$ | - | 2.5 | -300 | -300 | 0.022 | - | 0.10 | -610 | -600 |

**TABLE 2.** The values of the parameters of the Replicative Geometric, power-law, and exponential models as fitted to the speciation data of [15], to 2 significant figures. For each model the AIC and BIC values are reported, to 2 significant figures.

| Model | $\alpha$ | $\beta$ | $\gamma$ | AIC | BIC |
| --- | --- | --- | --- | --- | --- |
| Replicative Geometric | $1.3 \times 10^{-4}$ | 2.1 | 0.11 | -630 | -620 |
| Power-Law | $5.7 \times 10^{-4}$ | - | 1.4 | -570 | -570 |
| Exponential | $2.9 \times 10^{-3}$ | - | - | -120 | -120 |

**TABLE 3.**
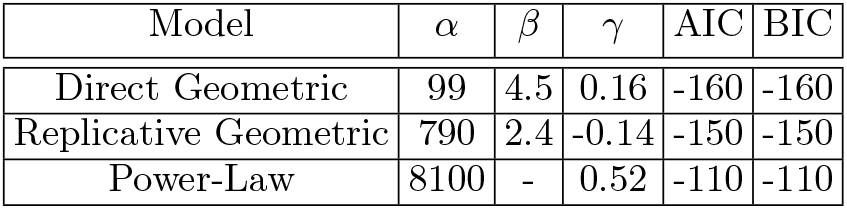
The values of the parameters of the Replicative Geometric, power-law, and Direct Geometric models as fitted to the arXiv submission data of [17], to 2 significant figures. For each model the AIC and BIC values are reported, to 2 significant figures.

**TABLE 4.** The values of the parameters of the Replicative Geometric, power-law, and Direct Geometric models as fitted to the patent submission data of [18], to 2 significant figures. For each model the AIC and BIC values are reported, to 2 significant figures.

| Model | $\alpha$ | $\beta$ | $\gamma$ | AIC | BIC |
| --- | --- | --- | --- | --- | --- |
| Direct Geometric | $1.1 \times 10^6$ | 0.13 | 0.57 | -170 | -160 |
| Power-Law | $5.1 \times 10^5$ | - | 0.55 | -110 | -110 |
| Replicative Geometric | $2.7 \times 10^{43}$ | -87 | $5.1 \times 10^{-3}$ | -110 | -110 |

**FIG. 1.**
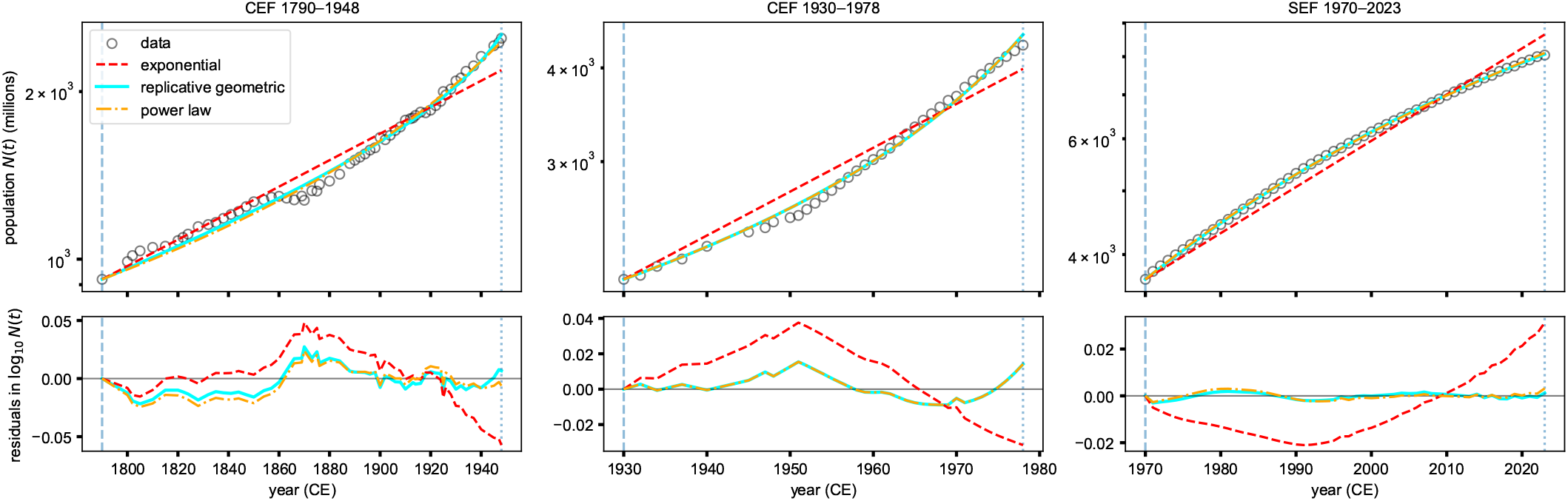
Fits of the Replicative Geometrical model, power-law growth, and exponential growth to three historical globalpopulation regimes using data from Ref. [12]. Within each interval the population is normalized according to Eq. 11. Left: 1790–1948, where the unconstrained Replicative Geometrical fit yields *γ ≃*2.2. Centre: 1930–1978, where the power law is slightly preferred and the Replicative Geometrical model approaches its small-*γ* power-law limit. Right: 1970–2023, where the Replicative Geometrical fit independently approaches *γ ≃* 1 with negative effective feedback.

**FIG. 2.**
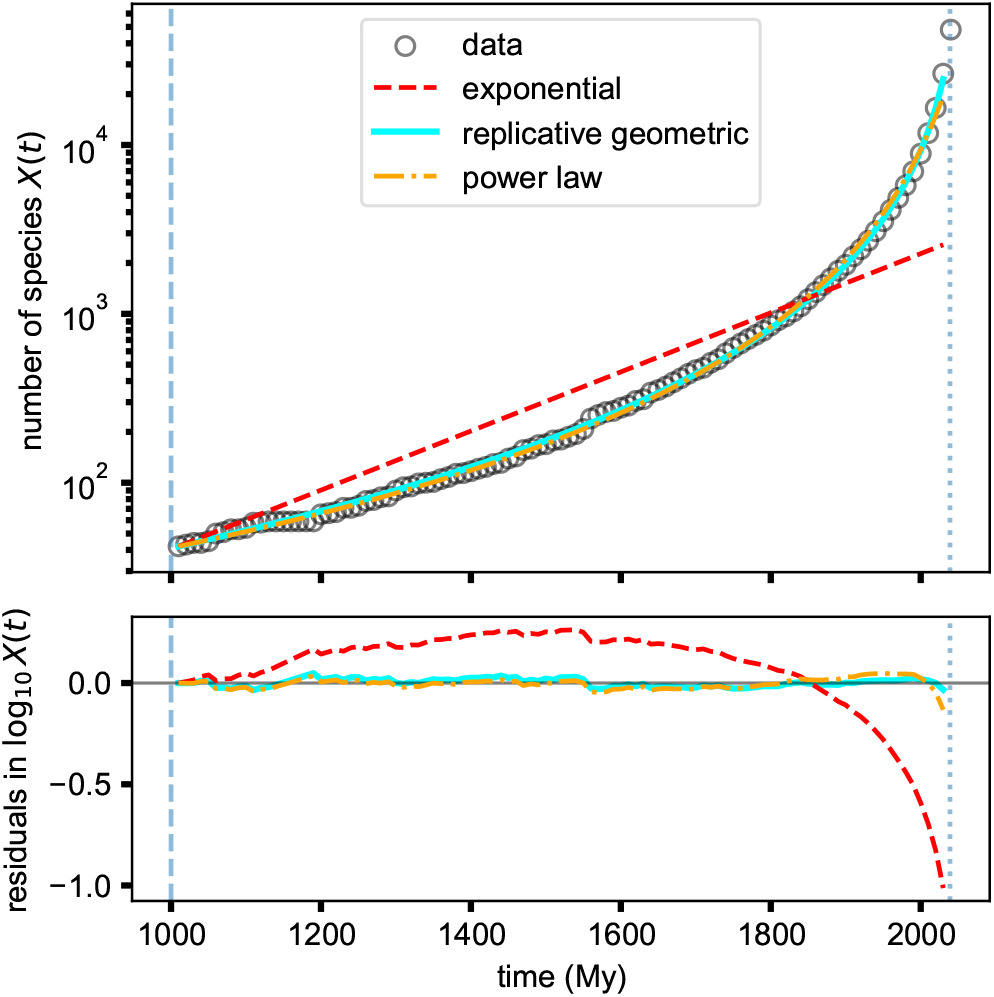
Comparison of the fitting of speciation data from [15] (circles), starting from *t* = 1000mY (vertical dashed line) to avoid low-sample bias and finishing 1mY before the end (vertical dotted line) to avoid selection bias, as suggested by Hedges *et al*.. The data is normalised to the initial value of the data-set, and sampled such that there is one data point every 10My to avoid over-sampling of data around *t ≈* 2000My.

For 1970–2023, the optimization instead yields *γ* ≃*s*1.0, with *α* ≃ 3.7 × 10^ℒ2^ yr^ℒ1^ and *β≃* ™0.57. It is notable that the original Trachenko–Zaccone exponent *γ* = 1 is recovered here without being imposed in the optimization. The negative value of *β* indicates that the net contribution of the effective component space becomes suppressive rather than growth-enhancing. Since *β* is a coarse-grained parameter, this observation does not by itself identify a unique microscopic demographic mechanism.

The 1930–1978 interval behaves differently. Here a power law is slightly preferred, with power-law exponent *p≃* 2.5, while the generalized fit is driven toward small *γ* and large *β*. This behaviour has a simple mathematical interpretation. For *γ ≪* 1,

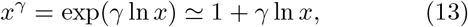

so that Eq. 12 becomes

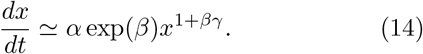

The power law is therefore an asymptotic boundary of the generalized geometrical family when *γ→* 0 while *βγ* remains finite. The 1930–1978 data are consistent with this limiting regime rather than with a well-resolved finite value of *γ*.

These fits suggest that the human-population record should not be represented by one fixed set of geometrical parameters over its entire history. Instead, the generalized law is naturally interpreted regime-wise: the topology and effectiveness of the social component space may change with historical conditions, allowing both the exponent *γ* and the effective feedback coefficient *β* to change between regimes. In particular, an effective *β <* 0 provides a mechanism for suppressed per-capita growth without requiring that the present paper introduce a separate logistic carrying-capacity factor.

Biological constraints nevertheless remain important. Equation 12 is intended as an effective law within a historical regime, not as an indefinitely valid asymptotic extrapolation. For *β >* 0 and *γ >* 0 the unconstrained equation exhibits finite-time divergence, whereas real populations can leave such a regime through changes in feedback, resource limitation, demographic transition, or other constraints. We therefore retain this point as a physical caveat rather than introducing a separate coupled carrying-capacity model here.

For completeness, an explicit bounded extension of the population dynamics is presented in Appendix A. This construction is not used in the empirical fits above, but demonstrates that a finite microscopic reproductive rate and carrying-capacity feedback are sufficient to regularize the unconstrained positive-feedback dynamics.

## V. POPULATION-DEPENDENT COMPONENT SPACES: SPECIATION

Speciation is another example where the individuals (species) contribute to the rise of new individuals. Let *X*(*t*) now denote the total number of species in a given environment. The rate of speciation, *dX/dt* ought to vary not only with the number of species (*dX/dt ~X*), but also with the number of available niches that can be exploited, which can be arrived at through mutation and selection. These niches may correspond to geographical (or, ‘abiotic’) features of the environment (and thus static, absorbed into the *dX/dt ~X* factor), but may also correspond to the presence of other species, or ‘biotic’ niches. Ecological opportunity arising from biotic interactions provides one route by which existing species can facilitate subsequent diversification [13]. The number of biotic niches can be imagined graphically, where each node corresponds to a species, and the number of edges per species, *E*(*t*) corresponds to the number of such niches. Through this lens, it has been empirically noted that the mean degree of this network evolves as *E*(*t*) *~X*(*t*)^*γ*^, where 0 *≤ γ ≤* 1 [14]. Under the assumption that biotic niches contribute independently to the rate of speciation, we obtain Eq. 10.

To explore the applicability of this model, we shall compare the fitting of contemporary models of speciation with data of speciation of eukaryotes [15]. We choose to compare the Replicative Geometric model with an exponential growth model, *dX/dt* = *αX* (which describes the birth-death model employed by Hedges *et al*.) and a power-law differential equation, *dX/dt* = *αX*^*γ*^, which could be an alternative interpretation of the rate of speciation varying with the number of biotic niches available to it.

Fig.2 demonstrates the accuracy of fitting to data which is sampled in 10mY intervals to avoid oversampling data in the *t ≈*2000My range, and Table. II demonstrates that the AIC and BIC values indicate the accuracy of the Replicative Geometric model is superior to the others considered.

## VI. POPULATION-DEPENDENT COMPONENT SPACES WITHOUT PER-CAPITA GROWTH: INNOVATION

Eq. 10 naturally describes human population and speciation because existing individuals and species necessarily give rise to other individuals or species, at a rate modulated independently by density of their relevant network structure, which scales as *X*^*γ*^. In addition to these examples, invention has also been analysed explicitly as a combinatorial process in arXiv citations and in U.S. patent data [16]. Unlike speciation or bacterial growth, we do not expect these markers of innovation to grow at a per-capita rate, as patents or manuscripts do not directly give rise to others in the same way bacteria give rise to more bacteria or species give rise to new species. Therefore, in these cases we anticipate a growth rate of the form

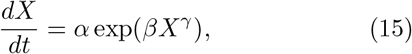

in other words, simply Eq. 10 without the *X* prefactor. As the individuals do not replicate, we term this a Direct Geometric growth model.

Using arXiv submissions and patent applications as empirical indicators of innovation, we compare the Direct Geometric model with the Replicative Geometric growth model and with the power-law form *dX/dt* = *αX*^*p*^. The latter provides an alternative description in which innovation production scales directly as a power of the existing innovation stock. As shown in Figs. 3 and 4, and quantified in Tables III and IV, the Direct Geometric model provides the best fit among the models considered for both the arXiv submission data [17] and the patent submission data [18].

**FIG. 3.**
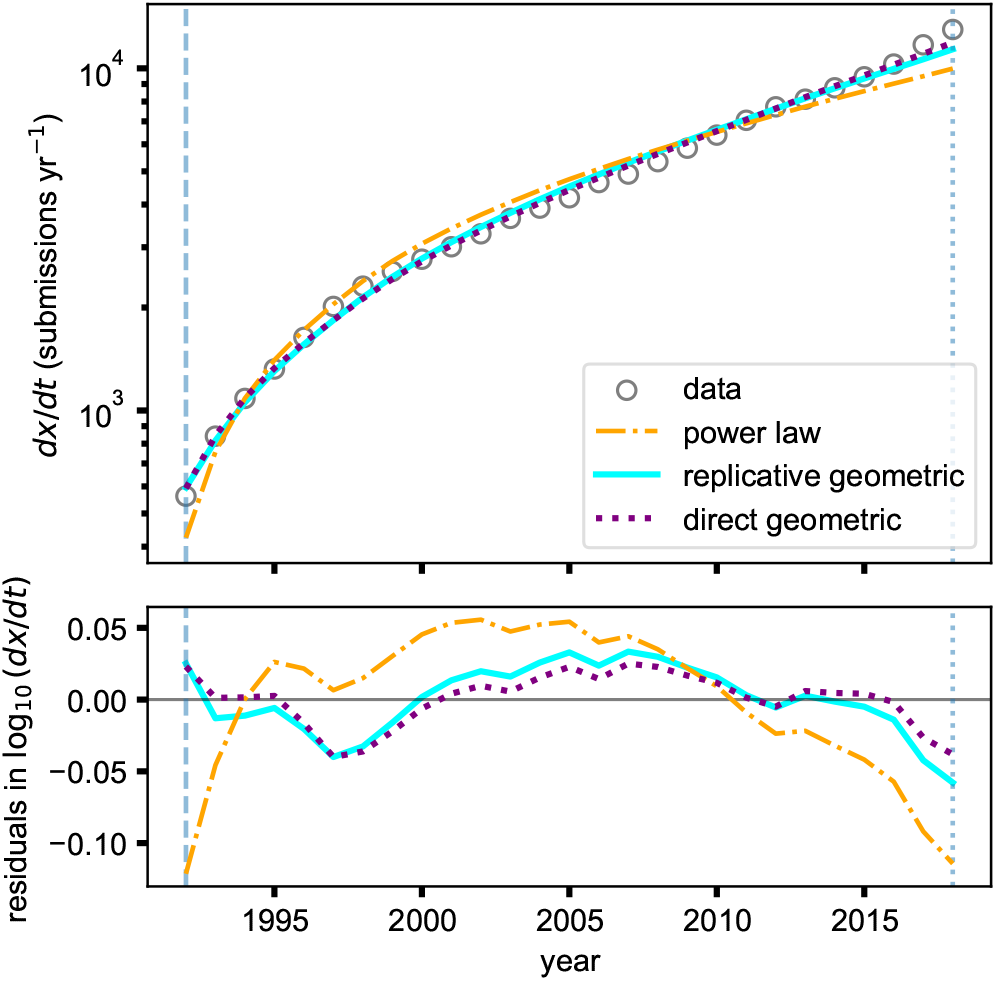
Comparison of the fitting of yearly arXiv submission data from [17] (circles), starting at 1992 (vertical dashed line) and finishing in 2019 (vertical dotted line). The number of new submissions per year is added cumulatively, *dX/dt* is drawn from this, and compared to three possible fits.

**FIG. 4.**
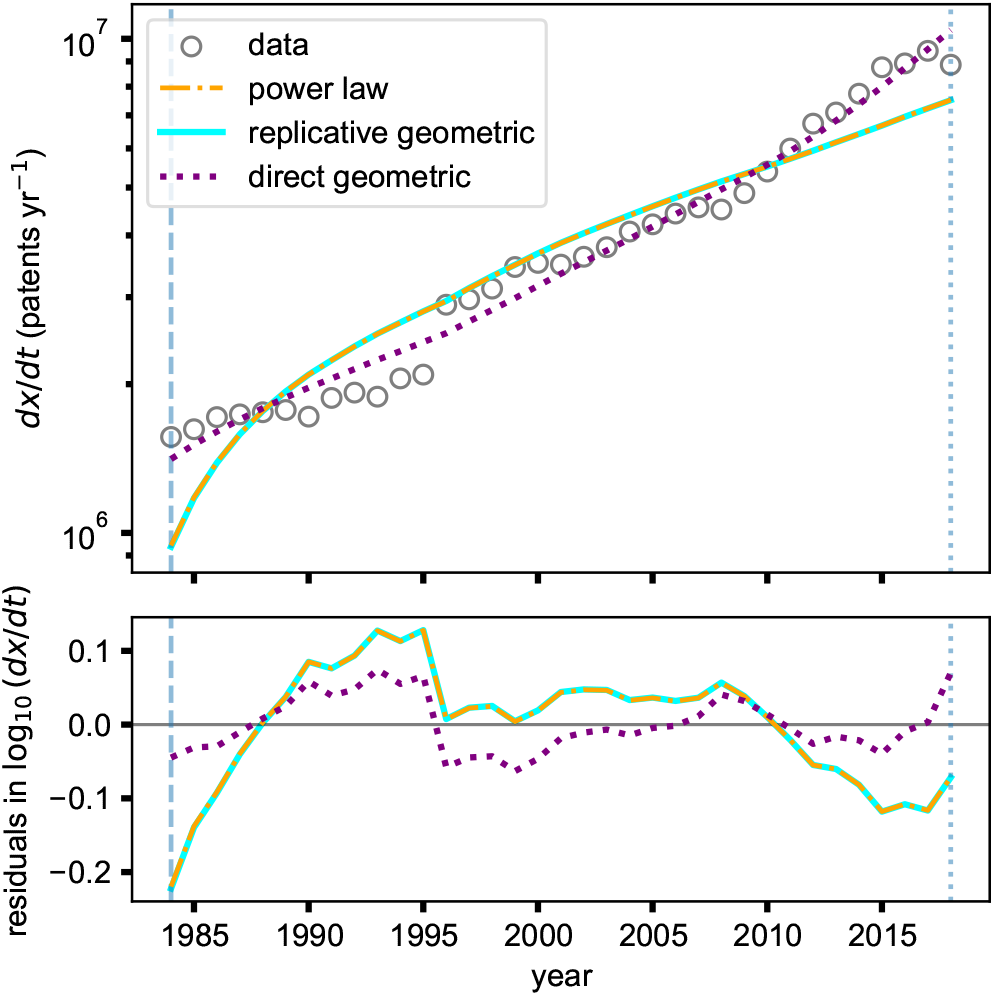
Comparison of the fitting of yearly patent submission data from [18] (circles), starting at 1984 (vertical dashed line) and finishing in 2019 (vertical dotted line). The number of new submissions per year is added cumulatively, *dX/dt* is drawn from this, and compared to three possible fits.

For the arXiv data, the Direct Geometric model yields lower AIC and BIC values than the Replicative Geometric model (Fig. 3). Moreover, the unconstrained Replicative Geometric fit converges to a negative value of *γ*, which is incompatible with its interpretation as an expanding effective component space. The Direct Geometric model therefore provides the more appropriate mechanistic description within the present framework.

The Direct Geometric model likewise yields lower AIC and BIC values than the Replicative Geometric model for the patent data (Fig. 4). In this case the Replicative Geometric fit converges to *β <* 0 together with *γ→* 0. A negative *β* corresponds to suppressive effective feedback, while the small-*γ* limit causes the model to approach a power law, analogously to the limiting behaviour discussed for the 1930–1978 human-population regime.

The model comparisons for each of these examples within the observed time windows are robust, but the mechanistic interpretation of the fitted parameters requires some caution. Neither dataset records the complete pre-existing stock of papers or patents that may contribute to the underlying interaction network before the beginning of the observational window. Consequently, the fitted values of the configuration-space parameters should be regarded as effective parameters for the recorded period rather than as universal network exponents.

## VII. DISCUSSION

The geometrical framework developed here identifies the size and activity of an underlying component space as two distinct determinants of growth. A common mathematical construction can therefore generate qualitatively different macroscopic growth laws depending on how these quantities evolve.

The bacterial example represents a fixed-size component space. The relevant physiological degrees of freedom are assumed to remain approximately fixed in number, while environmental conditions control their activation and deactivation. Growth dynamics is therefore governed primarily by the time dependence of *A*(*t*).

Human population growth, speciation, and innovation represent expanding-component-space regimes, distinct from the fixed-component-space bacterial case. Newly generated components may themselves enlarge the space of possible future configurations. When only combinations of fixed order are available, this produces power-law growth of configuration space. When combinations of progressively increasing order become accessible, exponential configuration-space growth becomes possible and naturally generates the generalised Trachenko–Zaccone nonlinear rate-feedback structure. We further distinguish classes of Replicative Geometric growth, in scenarios such as human population growth and speciation where individuals necessarily give rise to more individuals, and

Direct Geometric growth for cases where there is no percapita growth as in the case of markers of innovation like arxiv submissions or patent applications.

Human population growth provides a useful example of a component space whose effective feedback changes between historical regimes. The 1790–1948 interval yields a best-fit superlinear effective component-space exponent, whereas the 1970–2023 interval independently approaches the *γ* = 1 Trachenko–Zaccone limit but with negative feedback. The 1930–1978 interval instead approaches the power-law boundary of the same model family. These results favour a regime-wise interpretation of the geometrical feedback rather than a single fixed parameter set for the entire demographic record.

The generalized positive-feedback equation should therefore be interpreted as a description of a growth regime rather than an indefinitely valid asymptotic law. For positive feedback and *γ >* 0, the unconstrained equation can develop a finite-time divergence, whereas negative feedback suppresses the per-capita rate. Physical systems may leave a fitted regime through saturation, resource limitation, changes in network structure, changes in the sign of the effective feedback, or other constraints. The present work focuses on identifying the geometrical dependence within each regime rather than prescribing a universal saturation mechanism.

A major advantage of the present framework is that the competing growth mechanisms can be tested directly at the level of the per-capita rate for the Replicative geometric model. For a generic variable *X*,

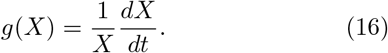

Constant-rate growth predicts

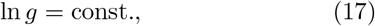

power-law feedback predicts

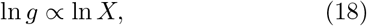

where as exponential rate feedback predicts

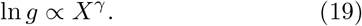

These transformed relations provide a transparent means of model discrimination that is complementary to direct nonlinear fitting of growth trajectories.

The same geometrical principle may thus connect growth phenomena across biological, demographic, technological, and ecological scales without requiring the microscopic mechanisms to be identical. What changes from one system to another is not the basic structure of the growth law, but the nature and evolution of the underlying component space that contributes to growth.

Future work should focus on determining empirically which class applies in particular systems. Bacterial growth data allow direct testing of the dependence of per-capita growth on active physiological components: while we have thus considered independent bacterial populations, synergistic or antagonistic [19] coculture of microbes may create a qualitatively different relevant component space. Future genomic analysis and fossil-record evidence will produce more data points to determine how the per-capita rate of speciation depends on local species richness, as opposed to the global species richness modelled here. For innovation, longer and more homogeneous patent and publication records can test the distinction between the Replicative and Direct Geometrical production laws. For human populations, demographic covariates may eventually allow changes in the effective feed-back coefficient to be linked to specific social or economic mechanisms rather than inferred only from population trajectories.

In this sense, the geometrical framework is intended as a common language for asking whether growth occurs within a fixed set of possibilities, or within a configuration space that is itself generated by the growth process.

## Appendix A

**A Bounded Extension of the Population Dynamics**

The Replicative Geometrical model describes the population dependence of the effective growth rate within a given regime. For positive feedback, however, an unconstrained component space can produce a finite-time divergence. A bounded biological continuation can be constructed by allowing the effective component-space size itself to saturate.

Let *M*_*b*_(*N, Z*) denote a bounded effective component-space size, where *Z*(*t*) represents environmental, technological, or social support conditions, and assume

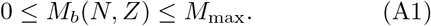

The corresponding geometrical per-capita growth rate is

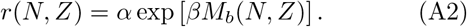

For *β >* 0, define

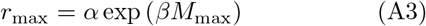

and

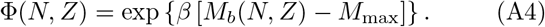

Since *M*_*b*_(*N, Z*) *≤ M*_max_,

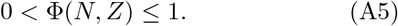

A carrying-capacity extension of the geometrical dynamics may then be written as

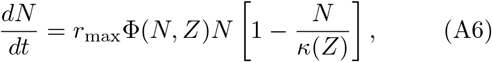

or, equivalently,

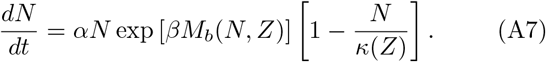

Thus the bounded equation retains the same geometrical rate factor as the main model; the additional assumptions are that the effective component space is bounded and that population growth may also be limited by an effective carrying capacity *κ*(*Z*) *>* 0.

### Theorem 1

*Let N* (*t*_0_) *be finite. If* 0 *<* Φ(*N, Z*) *≤*1 *and κ*(*Z*) *>* 0, *then on every finite interval over which Eq. A6 is defined, the population cannot exhibit a finite-time divergence*.

For *N ≥* 0,

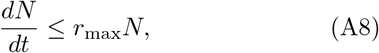

because Φ(*N, Z*) *≤* 1 and

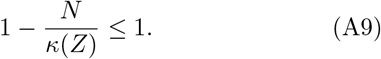

It follows that

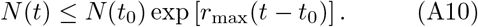

Hence *N* (*t*) remains finite at every finite time for which the model is defined.

Equation A6 is intended only as one possible continuation of the regime-wise geometrical dynamics studied in the main text. Other mechanisms, including changes in the sign or magnitude of the effective feedback coefficient, resource limitation, demographic transition, or changes in network structure, may equally cause a system to leave the unconstrained positive-feedback regime.

## Notes

https://github.com/tt386/GeometricalPopulationGrowth

### Competing Interest Statement

The authors have declared no competing interest.

